# Colors everywhere: enhanced chromatic processing across the first visual synapse in the zebrafish central brain

**DOI:** 10.1101/2020.06.19.160804

**Authors:** Drago A. Guggiana Nilo, Clemens Riegler, Mark Hübener, Florian Engert

## Abstract

Larval zebrafish (*Danio rerio*) are an ideal organism to study color vision, as their eye possesses four types of cone photoreceptors, covering most of the visible range and into the UV [1,2]. Additionally, their entire eye and nervous system are accessible to imaging, given they are naturally transparent [3–5]. Relying on this advantage, recent research has found that, through a set of color specific horizontal, bipolar and retinal ganglion cells (RGCs) [6–8], the eye then relays tetrachromatic information to several retino-recipient areas (RAs) [9,10]. The main RA is the optic tectum, receiving 97% of the RGC axons via the neuropil mass termed Arborization Field 10 (AF10) [11,12]. In this work, we aim to understand the processing of color signals at the interface between RGCs and their targets in the brain. We used 2-photon calcium imaging to separately measure the responses of RGCs and neurons in the dorsal brain to stimulation with four different colors in awake animals. We find that color information is widespread throughout the larval brain, with a large variety of color responses among RGCs, and an even greater diversity in their targets. Specific combinations of response types are localized to specific nuclei, but we observe no single color processing structure. In the main interface in this pathway, the connection between Arborization Field 10 and the tectum, we observe key elements of color processing such as enhanced signal decorrelation and improved decoding [13,14]. Finally, when presenting a richer set of stimuli, we identify parallel processing of color, motion and luminance information in the same cells/terminals, evidence of a rich color vision machinery in this small vertebrate brain.

## Stimulation with different colors evokes diverse responses in RGCs and central brain

Our goal is to characterize the flow of color information in the larval zebrafish brain, from the incoming RGCs and into the areas receiving these connections in the central brain. To that end, we used a custom-built microscope and 4-channel projector, capable of arbitrary color presentation (Fig. 1A-B, Supp. Fig. 1A) to deliver full-field, sinusoidal intensity modulated stimuli of four colors (Fig. 1C). The light sources were selected to broadly sample the range of wavelengths the fish is sensitive to at naturalistic intensities [6], rather than attempting to isolate and stimulate individual cone types [15]. Projection was from the bottom to stimulate the dorsal retina, which is well suited for color computations as it concentrates many color opponent RGC responses [8]. We utilized two transgenic fish lines, one expressing the fluorescent calcium indicator GCaMP6s [16] in RGC axon terminals, and the other expressing the same indicator in most of the larval central brain (Fig. 1D).

**Figure 1.**
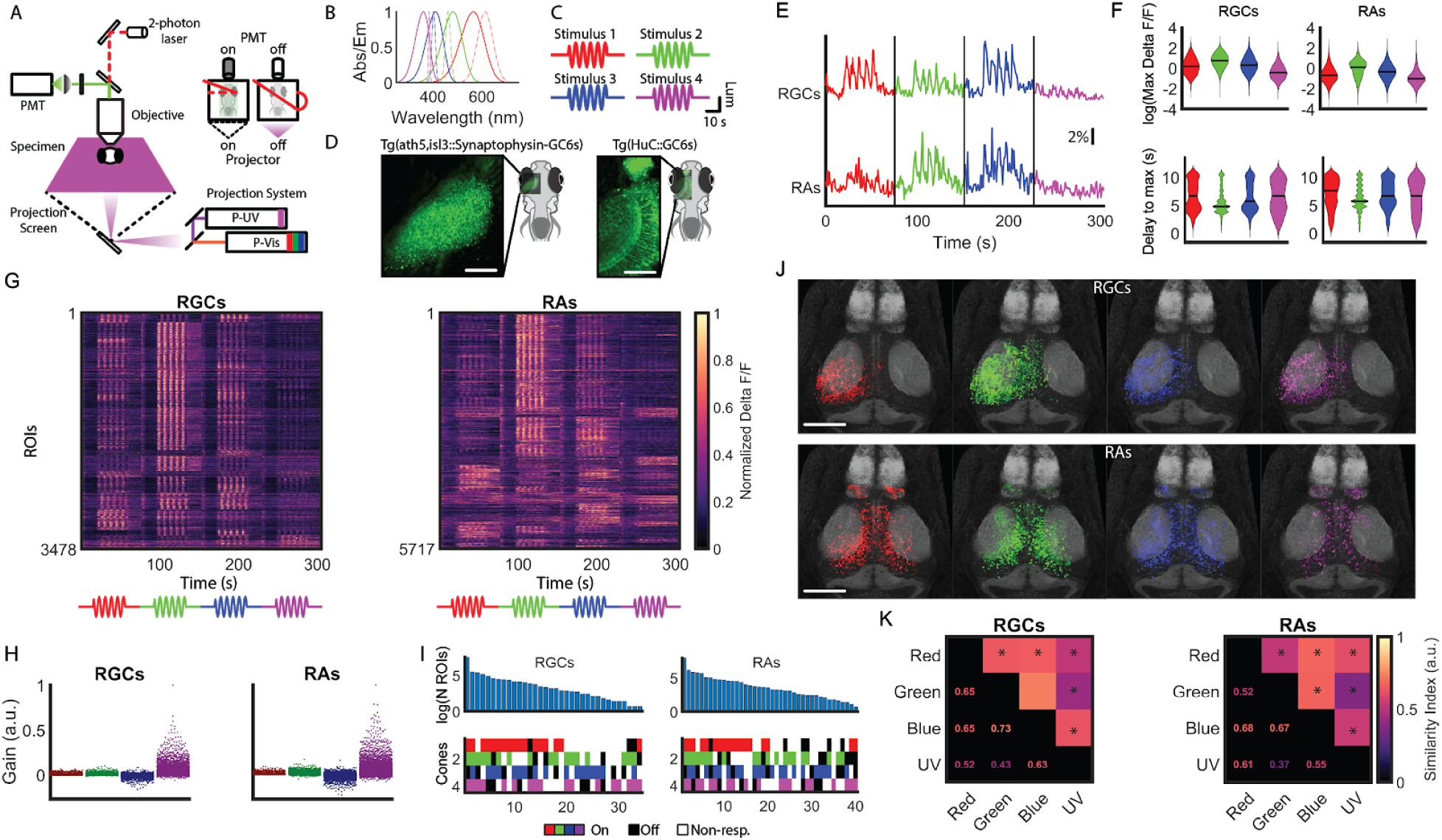
Stimulation with different colors evokes a variety of responses in zebrafish RGCs and RAs. A: schematic of the 2-photon microscope used for stimulation, projecting color stimuli from the bottom using a custom-designed DLP projector with four channels. (inset) alternation between stimulation and recording during the mirror turning time. B: absorption spectra of the zebrafish cones (dotted lines) and emission spectra of the LEDs (solid lines) used in the projector. C: full field, temporally sinusoidal stimuli utilized. D: average images from the isl3 and HuC:GCaMP6s lines utilized to label RGCs and RAs respectively. E: representative delta F/F responses to all 4 stimuli from RGCs and RAs. F: maximum delta F/F (top) and delay to peak response (bottom) for the response to each stimulus in RGCs and RAs. G: full datasets for RGCs and RAs, with intensity reflecting the row-normalized delta F/F. H: calculated gains for each cone from each ROI in both datasets. I: response types obtained from categorizing the different combinations of cone gains from both RGCs and RAs. J: registered anatomical maps of the ROIs, colored by the stimulus eliciting the maximum response in each ROI. K: similarity of the response patterns elicited by the four color stimuli. A value close to 1 indicates identical patterns, and a value close to 0 indicates perfectly non-overlapping patterns. Star denotes overlap significantly different from inter-trial overlap, Wilcoxon signed rank, p<0.05. For all panels, N=6 animals and 3478 ROIs for the RGCs data and N=6 animals and 5717 ROIs for the RA data. Scale bar in D: 70 μm. Scale bar in J: 90 μm.

We found that all four color stimuli elicited responses in multiple brain regions (Fig. 1E). We observed similar overall response maxima and delays to peak for all four stimulation colors in both the RGCs and RAs, although the responses were slightly slower and weaker in RAs (Fig. 1F). We recorded in total 3478 ROIs (Regions of Interest) from six fish in RGCs and 5717 ROIs from six fish in RAs. We define ROIs as small clusters of terminals or cells isolated via correlated activity, see Methods for details [4]. Most of the responses were dominated by the green stimulus, although there were many other patterns (Fig. 1G). Also, the responses in the RAs were more color specific than the responses in RGCs, with a tendency for one or two color stimuli to dominate the response.

To better probe the variability of the responses, we constructed a linear model to infer the required cone type’s gains (amount of response contribution) yielding a given ROI response [15,17]. As has been reported before, gains of the UV cones were considerably higher than those of the other cone types, and blue gains were mostly negative, indicating they have Off polarity (Fig. 1H and Supp. Fig. 1B-C, [8,18–20]). The distribution of the gains of RGCs and RAs were very similar, so we identified the patterns present in the combinations of gains to probe for potential differences. To simplify the identification, we discretized the gains as being positive, negative or 0. Indeed, the RAs showed a larger number of response types than RGCs (Fig. 1I). Most of the patterns showed some degree of color opponency, but as reported before, far fewer than the possible 81 combinations (four cones, each with on/off/non-responsive states, i.e. 3^4^ different options) were actually present, possibly due to ethological reasons such as the spectral composition of natural light [6].

We wondered whether the diverse responses we observed have a particular anatomical distribution, so we registered all the responses in all the animals to a reference brain [21] and visualized their location on the optic tract and the brain. These distributions showed partial color specific spatial organization (Fig. 1J). We quantified this as the amount of overlap of the patterns of activity elicited by each color, normalized by the overlap between the patterns evoked in single trials for the same color (Fig. 1K). The similarity was lower in RAs than in RGCs, although this could be due to the anatomical constraints of the optic tract. We also observed that similarity was lower for UV than for the other colors in RGCs, but this levels out in the RAs.

## Color responses cluster functionally and anatomically

Given the observed localization, we next assessed whether different brain regions within the RGC AFs and RAs displayed selective types of color responses or color processing (Fig. 2A-B). The different regions contain diverse response types based on their averages (Fig. 2C). Among the AFs, AF8 shows a similar average response as AF10, mostly of the On type, but AF9 and AF4 contain a mixture of On and Off responses, and AF5 has a less green dominated average response. Variability is also observed across RAs, where the tectum and pretectum show faster responses, while the cerebellum and habenula show non-oscillating, sustained signals (Supp. Fig. 2A-B).

**Figure 2.**
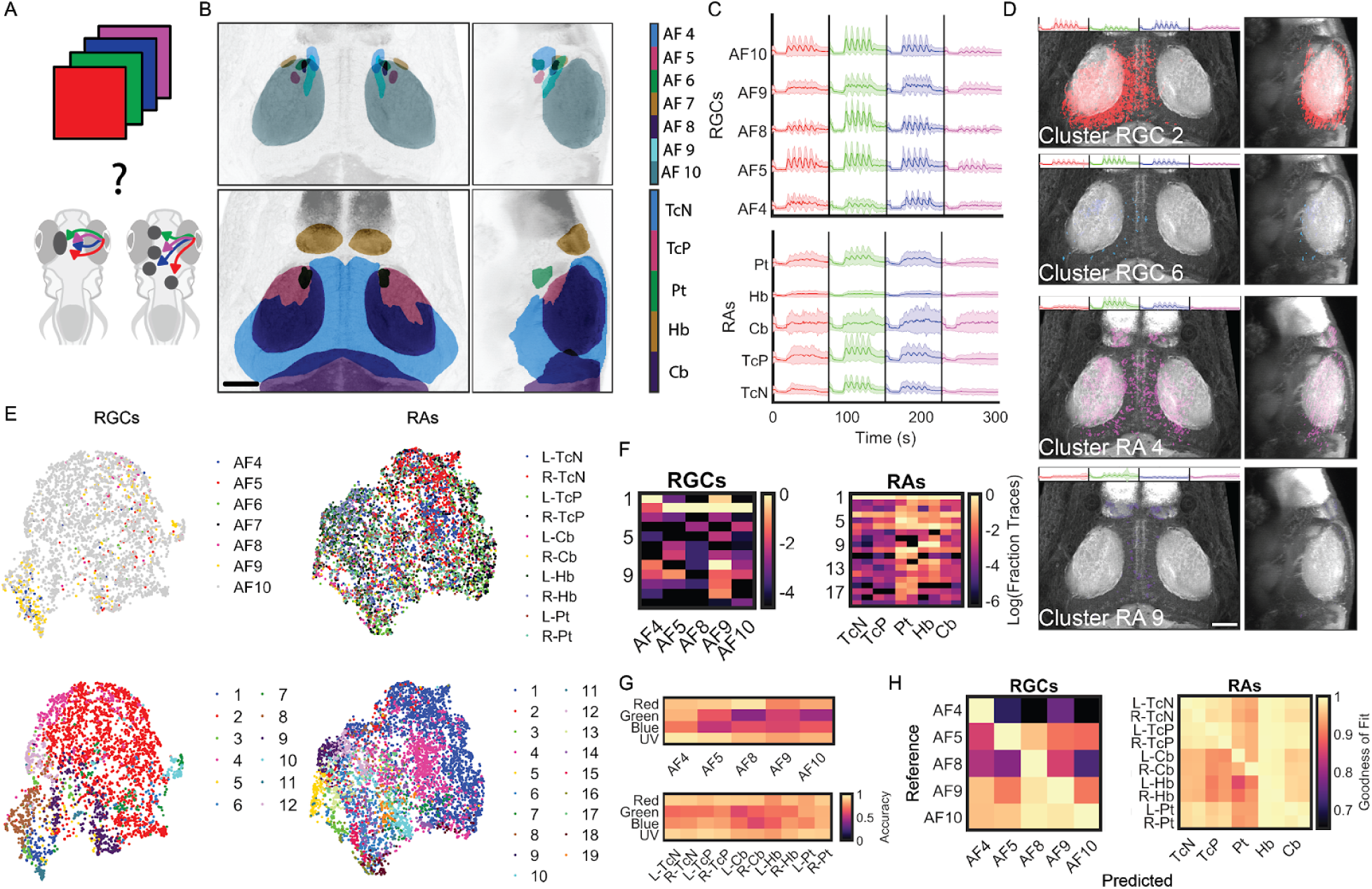
Color responses tend to cluster functionally and anatomically. A: we presented four color stimuli and measured responses from a number of brain regions to identify color-processing areas. B: anatomical location of the brain regions imaged within the RGC and RA animals. C: average delta F/F traces for each stimulus from each of the regions recorded in both strains. D: registered locations of four example clusters in the reference brain, two for the RGC and two for the RA datasets. Inset shows the average delta F/F traces for these functional clusters. E: UMAP embedding of the ROIs for RGCs and RAs, using their respective Principal Component decompositions as the source for the embedding.They were colored according to the region of origin of the ROI (top) or the functional cluster each ROI belongs to (bottom). F: relative proportion of each cluster found in each of the brain regions imaged (normalized per brain region). G: average performance of an SVM classifier trained to discriminate between the different colored stimuli on a per-region basis for the RGC and RA brain regions. H: average r-squared coefficient for each pair of regions when using the clusters of one region to approximate the ones in the other. Shown for RGCs and RAs. For RAs in F and H, the left and right hemispheres of each brain region correspond to the left and right column respectively. For all panels, N=6 animals and 3478 ROIs for the RA data and N=6 animals and 5717 ROIs for the RGC data. Scale bar in all maps is 60 μm. AF: arborization field, R-, L- : right or left hemisphere, TcN: tectal neuropil region, TcP: tectal periventricular region, Cb: cerebellum, Hb: habenula, Pt: pretectum.

To better understand this variability in the responses, we utilized Gaussian Mixture Model-based clustering to group the responses by similarity (Supp. Fig. 2C-D, [22]). We observed a larger variety of responses in the RA dataset, encompassing different combinations of intensities and polarities (On/Off) for each stimulus. Additionally, these clusters were well distributed across animals in number and type (Supp. Fig. 2E-F). Are these clusters differentially associated with different brain regions? To address this question, we used registration to map the cluster locations onto the reference brain utilized in Figure 1J. Figure 2D shows two example clusters from each RGCs and RAs, illustrating that they show partial regionalization.

To obtain a whole-dataset view of this grouping, we employed UMAP (Uniform Manifold Approximation and Projection, Fig. 2E [23]). Briefly, we used the response features of each ROI to embed it in a reduced dimensionality space, and then colored each data point according to either the region (top), or the cluster (bottom) corresponding to that ROI. Using this approach we find considerable overlap between the clusters and regions. We quantified this by evaluating which proportion of the responses in each region corresponded to each cluster (Fig. 2F). Most regions were dominated by one or two clusters, with the observed layout recapitulating described connectivity [11]. Namely, AF4 and AF9 share functional patterns, and it is known that these regions receive a subpopulation of RGC projections, which do not terminate in AF10. The other areas are more similar to AF10, but they are also enriched in other clusters. In the RAs, the cerebellum and habenula separate themselves from the tectal response pattern, and so does the pretectum to a lesser extent, again recapitulating what is known about retino-recipient targets [9]. In summary, although there is no obvious unique color sensing area in the imaged brain regions, there is a distinct, area specific pattern of color response types, both in RGCs and RAs, suggesting that the responses identified through these clusters are used for different color computations.

We next assessed how well an ideal observer could distinguish colors from the responses in each region. To this end, we trained classifiers (Support Vector Machines, subsampled to match ROI number) on the responses from each animal to understand which components of the stimulus were being encoded. Although the decoding capacity of most areas is quite high, the two colors in the middle of the spectrum (green and blue) are separated less accurately (Fig. 2G, middle rows). Additionally, AF4 seems to outperform even the much larger AF10 in the classification (Supp. Fig. 2G). The RAs also performed well, with the cerebellum being the worst performing relative to the others, and the pretectum slightly outperforming the tectum. Across all regions, green and blue were the most difficult stimuli to distinguish. Since distinguishing stimuli could be based on the extraction of pure intensity information from the incoming light, we also trained an SVM to classify the individual intensity levels for each stimulus separately (Supp. Fig. 2H). While the performance is above chance, it appears information integration beyond intensity levels is required to classify colors. Taken together, colors were well distinguished across brain regions, but the observed differences suggest a degree of region-specific processing.

To corroborate these results, we used a regression approach to fit every cluster in an area with the clusters in the other (Fig. 2H). These results confirm that most color information is present in AF10 in the RGC group, and that there is common information between AF4 and AF9, given their shared goodness of fit. In RAs, again cerebellum underperforms, but the other areas are very balanced, despite the difference in their response profiles.

## Colors are better separated in the tectum than in AF10

AF10 and the tectum are the largest regions for visual information processing in fish, and the former directly follows the latter in the visual pathway. We therefore sought to better understand the processing of color signals across this connection (Fig. 3A). We find that the tectum surpasses AF10 in color classification (Fig. 2G, Supp. Fig. 3A-B), even though they are separated by only one synapse. To analyse this further, we focused first on the single ROI level using UMAP to search for differences in the responses to particular stimuli. For this, we ranked all ROIs based on their response amplitude to each stimulus. We next assigned colors based on which ROI belonged to the top 75^th^ percentile of each stimulus, or their combination, when ranked based on evoked response (Fig. 3B). It becomes evident that many ROIs are not in the top responders of any stimulus (Fig. 3C), and that there are about equal proportions of responders to each color or their combinations between AF10 and tectum (Wilcoxon signed rank test, p>0.05). This is also observed if we instead look at the cone gains.

**Figure 3.**
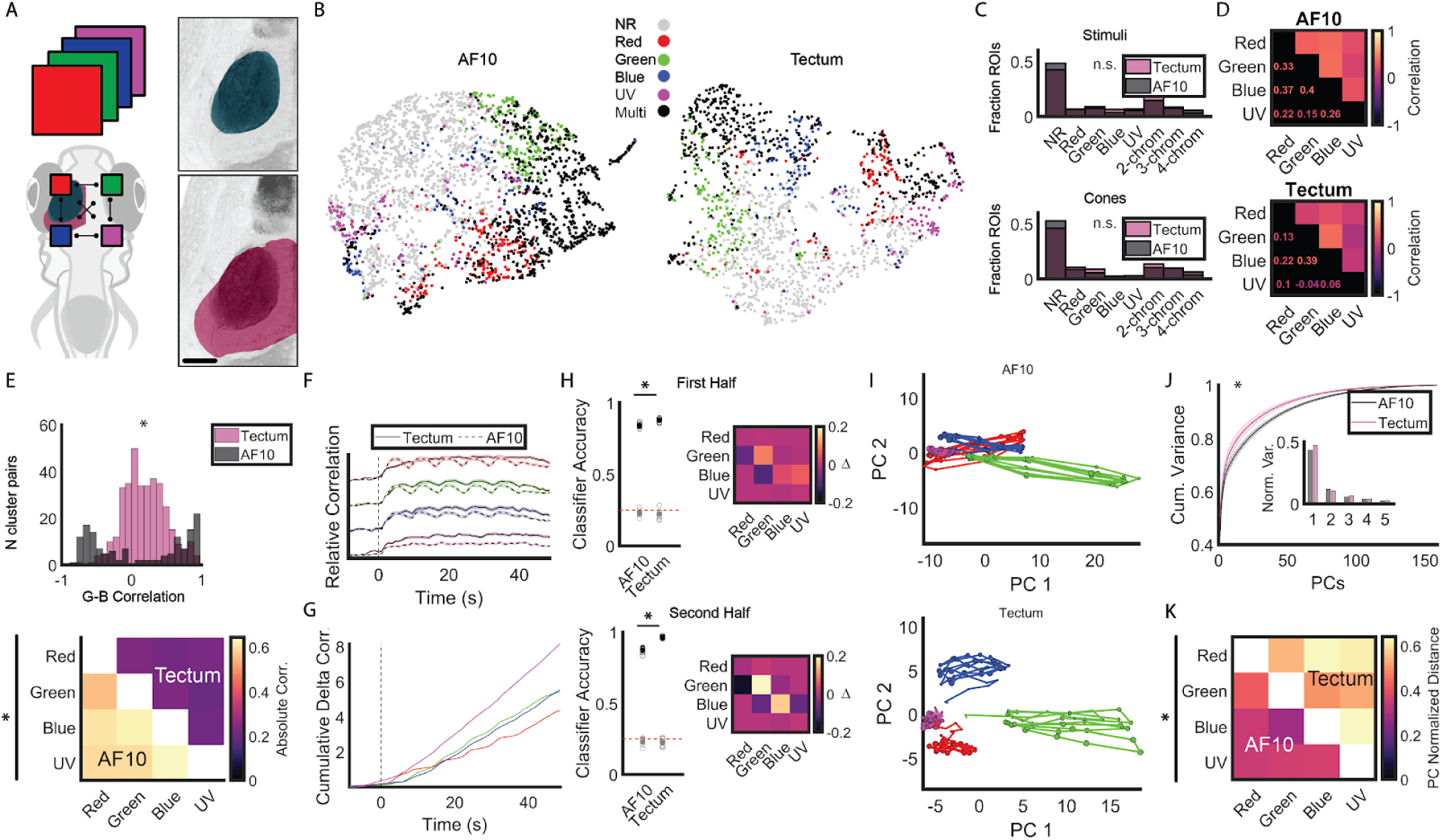
Colors are better separated in the tectum than in RGCs. A: (left) the same data as before was utilized to assess the color computations occurring between AF10 and tectum. (right) anatomical layout of AF10 (top) and ipsilateral tectum (bottom), used for this section. B: UMAP embedding of the principal components for each dataset, colored according to the ROIs belonging to the top 75th percentile of responders for any given stimulus. Gray circles denote non-responsive (NR) ROIs and black circles denote ROIs assigned to more than one stimulus. C: (top) histogram quantifying the number of ROIs assigned to the top 75th percentile of each stimulus, no stimulus or multiple stimuli for both groups of fish. p>0.05 for all bins tested separately, Wilcoxon signed rank. (bottom) same as in C for the cone gains. D: overall correlation of the neural activity evoked by each stimulus for each dataset. E: (top) cluster average correlations between the green and blue stimuli for the tectum and AF10. (bottom) cluster average correlations for all pairs of stimuli for AF10 (below diagonal) and tectum (above diagonal). The deltas were evaluated pairwise, p<0.05 Wilcoxon rank sum. F: within-stimulus correlation in the neural population over time at each frame for AF10 and tectum. The dotted line indicates the stimulus onset. G: cumulative delta over time between AF10 and tectum for the data shown in F. H: SVM classifier trained to discriminate between the four colored stimuli using the responses from either the first (top) or the second half of the stimulation period (bottom). The black filled circle represents the average of the individual classification repetitions, shown in black open circles. The gray filled circle is the average of the shuffled repetitions, shown in open gray circles. Star denotes p<0.05, two-tailed Wilcoxon signed rank test between the non-shuffle data. (right) Delta confusion matrices for both classifiers, showing the difference in individual stimulus performances between the tectum and AF10. Presented stimuli are on the rows and predicted stimuli on the columns. The red line indicates performance with shuffled labels. I: CCA-aligned and averaged Principal Component trajectories over the course of the stimulation period for AF10 and tectum. The color of the line indicates the stimulus and the size of the dot increases over time. J: cumulative principal component variances and s.e.m. for the tectum and AF10 across all animals. Star denotes p<0.05, Wilcoxon rank sum. (inset) Relative variances of the first 5 PCs for AF10 and tectum. K: average pairwise euclidean distances between the centroids of the point clouds corresponding to each stimulus responses in PC space for AF10 (below diagonal) and tectum (above diagonal) after alignment. Star denotes p<0.05 in a Wilcoxon rank sum test performed on the corresponding pairs of stimuli. For panels A-H and J, N=6 animals and 1776 ROIs for the tectum data and N=6 animals and 2670 ROIs for the AF10 data. For panels I and K, N=3 animals and 1604 ROIs for the tectum and N=5 and 1313 ROIs for AF10. Scale bar in A is 60 μm.

We next wondered how the responses contribute to computation within the population. Since it has been shown that pattern decorrelation is a hallmark of sensory neural computations [13], we computed the correlation of the responses between each pair of stimuli for all ROIs in AF10 and tectum (Fig. 3D, Supp. Fig. 3C). We found that the population responses are indeed less well correlated in tectum than in AF10, but the difference is small, especially for the green-blue pair. As this overall correlation collapses a large amount of information, we then tested whether correlation between the functional cluster average responses could reveal processing differences (Fig. 3E). Indeed, for every pair of colors (with green-blue as an example on top), the correlations between averages for each stimulus pair are lower for tectum than for AF10, which tend to be either very high or very low (p<0.05, Wilcoxon rank sum). This begs the question of how the responses to each color are differently represented in AF10 vs the tectum. To address this, we first focused on the temporal structure of the response. We found that the correlation between the activity of each pair of neurons (Fig. 3F, Supp. Fig. 3D), in response to all four colors, evolves over time in the tectum, while AF10 remains stably oscillating with the stimulus. The cumulative delta between AF10 and tectum (Fig. 3G) shows clearly how, after stimulus onset, the tectum responses diverge from responses in AF10, which suggests that time does play a role in the long term evolution of the response, potentially for adaptation purposes. To put this time evolution to the test, we used an SVM classifier trained on either the first or the second half of the stimulus period to decode the color of the stimulus (Fig. 3H). While both the tectum and AF10 performances are balanced in the first half of the period, the tectum outperforms AF10 by ∼10% when trained on the second half, supporting our hypothesis.

To gain further insight into the activity patterns leading to the observed decorrelation, we projected the neural responses into a lower dimensional space using CCA-aligned PCA (Canonical Correlation Analysis, [24] and see methods). We observed that AF10 shows activity patterns that closely follow the stimulus sinusoidal intensity fluctuations (Fig. 3I, Supp. Fig. 3E-F). In addition, the trajectories evoked by the four stimuli mostly traversed space along the axis of principal component 1. In the tectum, in contrast, the trajectories evolved over time in a divergent pattern, where every color evolved in a different direction. Therefore, apart from time, there is a dimensional component to the improved separation. We first quantified this difference as the weight distribution across principal components (Fig. 3J). For most components except the second, the tectal principal components explain more relative variance than their AF10 counterparts (p<0.05, Wilcoxon rank sum). We also measured the relative distances between the centroids of the spaces occupied by each stimulus (Fig. 3K). The distances between the point clouds in the tectum are greater than the distances in AF10 for all stimulus combinations (p<0.05, Wilcoxon rank sum). Overall, this shows that the tectum surpasses AF10 in color processing capabilities due mainly to a combination of time evolution and higher dimensionality.

## Color separation extends to different spatiotemporal patterns

So far, the color stimuli we utilized contained mainly chromatic information, but the visual world of many organisms contains parallel layers of information that include not only color, but also local contrast, movement and shapes [25]. Therefore, we employed another set of stimuli that include the aforementioned properties, namely a checkerboard oscillating in luminance, a moving grating, and a flat intensity background with a dark flash in the middle of the stimulation period (Fig. 4A-B). These were presented in red and UV to the tectum and AF10 as above, as it has been suggested that many color comparisons in the retina occur between high and low wavelength illumination [26], and since these two colors stimulate non-overlapping sets of cones (Fig. 1B). On average, the responses elicited by all stimuli were very similar between AF10 and tectum (see also Supp. Fig. 4), although the red grating evoked a higher response in the tectum. This could be explained by the high proportion of direction selective cells in this structure, but it does not explain why this is not the case for UV (Fig. 4C, [27,28]). Inspection of the trial averaged responses reveals a higher diversity when compared to the four sinusoidal color stimuli, given the increased complexity of the stimuli, although this is more obvious in tectum than in AF10 (Fig. 4D).

**Figure 4.**
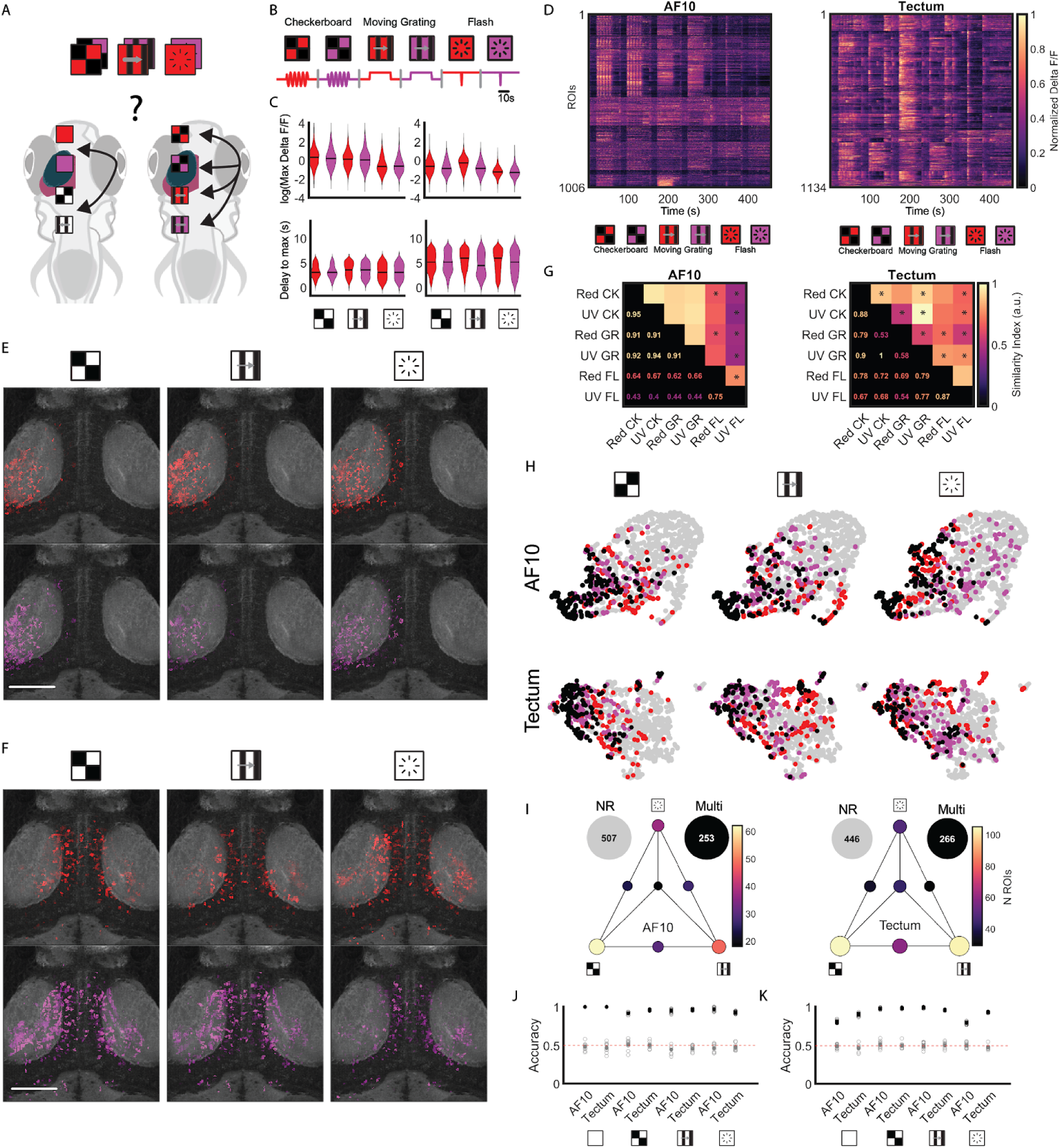
Color separation extends to different spatiotemporal patterns. A: an additional set of stimuli was utilized to assess whether different spatiotemporal patterns affect computation of color between AF10 and tectum. B: diagram of the stimuli utilized, combining spatiotemporal patterns with red and UV. C: maximum response (top) or delay to peak (bottom) for each stimulus and brain region. D: trial averaged fluorescence traces. Intensity is row-normalized fluorescence. E: anatomical ROI distribution from the AF10 terminals, depicted according to the stimulus that elicited the maximum response for that ROI. F: same as in E for the tectum. G: anatomical similarity matrix for the response patterns elicited by each stimulus. Star denotes overlap significantly different from inter-trial overlap, Wilcoxon signed rank, p<0.05. H: UMAP embedding of the ROIs based on their principal components. Color was assigned based on whether an ROI belonged to the 75th percentile of responders for the indicated stimuli, for both the AF10 (top) and tectum (bottom) ROIs. The color indicates the stimulus an ROI belongs to, with gray indicating non-responders and black indicating the ROI responded to both stimuli. I: Graph quantifying the distribution of single color sensitive ROIs (only in the top 75th percentile for one color). The symbols indicate the nodes corresponding to only that pattern, while the nodes in between correspond to ROIs selective for that combination of patterns. The color and size of the node indicate the number of ROIs. The outside nodes indicate ROIs not included in the graph: NR: ROIs not present in any set. Multi: ROIs sensitive to both colors. J: classification performance of an SVM classifier trained to determine the color of the stimulus for each one of the spatiotemporal patterns, including the full field stimulus performance for comparison. The filled black circle is the mean, the open black circles the individual repetitions, the filled gray circle is the average of the shuffled labels classifier and the open gray circles are the individual shuffle repetitions. K: same as in I for a classifier trained to discern between the pre-stimulus period and the stimulus period for both colors. The red line indicates random performance. For all panels N = 4 animals, 1006 ROIs for the AF10 dataset, N = 11 animals, 1134 ROIs for the Tectal dataset. Scale bar in E-F 90 μm. CL: checkerboard, GR: moving grating, FL: dark flash.

Anatomical registration onto the reference brain reveals distinct patterns of activation in AF10 and in tectum in the registered data, both in the color and spatiotemporal pattern dimension (Fig. 4E-F). These patterns are more similar in AF10, especially when comparing between the checkerboard and gratings for each color channel. When calculating the pattern similarity, the AF10 patterns indeed are more similar for the non-flash stimuli, while in the tectum the overall similarity decreases (Fig. 4G).

Given the high anatomical overlap of these populations, we tested whether this is preserved functionally at the ROI level. We projected the ROIs into 2D space using UMAP as before, and visualized all responses lying in the top 75^th^ percentile for each stimulus (Fig. 4H). The patterns in tectum are more widespread and specific, i.e. fewer ROIs respond to both colors, especially for the moving gratings and the flash. When visualizing the preferred stimulus for the ROIs that responded to only one color (∼50% of the total ROIs in the top 75^th^ percentile), it is evident that they are distributed throughout the stimuli, although the ROIs that were in the top responders for only one stimulus are more common than the ones sensitive to 2 or 3 stimuli, especially in tectum (Fig. 4I, see lighter colors on the vertices of the triangle graph).

Given this specificity, we next tested whether it was possible to decode stimulus information from the responses. As before, we trained an SVM to distinguish between the color of the stimulus, and trained it separately on each of the spatiotemporal patterns, including, for comparison, training on the pure color stimulus from the previous stimulus set (Fig. 4J). The neuronal populations were able to decode color at comparable performance across pattern and brain region. Since color information is preserved, we wondered whether the stimulus information was present, or whether we were mostly recording from color-specific cells. For testing this, we trained an SVM to discern between the pre-stimulus and stimulus period regardless of color, as this should isolate decoding of the contrast changes, motion and flash features by the different spatiotemporal patterns (Fig. 4K). Indeed, the SVM performed above chance for all the stimuli, indicating that by and large we recorded from a mixed population, and that the information about color seems to be intermingled with the information about other visual modalities, at least at the level of the tectum.

## Discussion

In this work, we show how color stimuli are processed in two successive stages of the larval zebrafish visual pathway. We found that color information is widely available across the larval brain, even in cerebellum and habenula, which are not usually associated with color processing in other vertebrates [29–31]. While the incoming RGCs and the retino-recipient areas we surveyed carry in general common sets of information, the particular response types in each area differed, and therefore offer varied computational capabilities. When focusing on AF10 and the optic tectum, we observed that the color computations are more advanced in tectum, as evidenced by improved decoding and better pattern decorrelation over time, while the responses in AF10 followed the stimulus fluctuations more closely. Finally, we compared the color responses to different behaviorally relevant stimuli, and found that color information is available in conjunction with other visual features, speaking in favor of its influence in many zebrafish brain processes.

As this is the first report of color processing in the zebrafish central brain [8], our aim was to characterize its anatomical distribution. RGCs are known to arborize in defined areas of the brain, and there is cumulative evidence pointing to the selectivity of these areas for certain types of responses [32,33]. This seems to be also the case with color information. Despite AF10 receiving the majority of the connections from the retina, AF4 and AF9 seem to receive different response types than AF10. This is consistent with previous reports of the retinal projectome, indicating that AF9 receives inputs that do not continue to AF10 [11]. In particular, both AF4 and AF9 receive a strongly Off-biased response complement, which is indeed reflected in the averaged cluster responses of these areas (Supp. Fig 2C, Cluster 8)

Likewise, although the optic tectum dominates the diversity of response types, other areas like the cerebellum and the habenula also show very unique response profiles, as indicated by their particular collection of clusters, such as a prevalence of non-oscillating, sustained responses (Cluster 11 in cerebellum). This could be evidence of further response processing, even related to the generation of premotor signals given the known roles of this brain region [34]. Additionally, the reported lateralization of habenular responses [33,35] is also reflected at the level of color, as the left habenula seems to have an enrichment of green evoked responses while the right habenula receives a wider response complement. ROIs in these brain areas are also stimulated mostly by a single color, which is not predominant in other brain regions. Further studies will need to address whether the presence of color information in any of these brain regions has a causal role in visual processing.

Recently, Zhou et al. [8] described the spectral properties of RGC dendrites and somata in the eye of the larval zebrafish. Our results corroborate their finding of mostly Off blue weights and high UV weights in RGCs, and also of simple and complex color opponencies, although the proportions of types we observed are different. This could be due to a number of factors, including differences in the amount of innervation from each RGC type to the central brain, the difference in stimulation protocols and type calculation, and our correlation-based ROI isolation in a densely labeled preparation. Interestingly, Zhou et al. also describe the presence of a time code in RGC responses. Due to our imaging protocol, we are unable to resolve the calcium dynamics at the rates this time code is observed, but in future studies, it will be interesting to elucidate the interaction between the slower time component we observe with the described RGC time code and color processing.

In mammals, and in particular in primates, color information follows a processing path down the temporal stream, from V1 to V2 and V4 [36]. In the latter, one can find neural correlates of more complex aspects such as hue. Overall, we observed no obvious color processing center in the fish brain, as color responses appear anatomically distributed. This does not preclude its existence of course, as future studies with an increased phase space of exploration (such as the multiple frequency sequences in [8]) and more anatomically restricted imaging might be able to find such a region.

While the exact role of color vision in the larval zebrafish life cycle is only starting to be understood, it is encouraging that chromatic information is widespread throughout the brain, even in common channels with other essential components of vision such as luminance and motion. Future studies should be able to parse this color richness, and go deeper into our understanding of color computations throughout the whole brain, as well as the parallel processing of these computations together with other visual modalities, such as the processing of contrasts and spatial patterns.

## Acknowledgments

The authors would like to thank Martin Haesemeyer, Owen Randlett and Tom Baden for their contributions to data analysis, and Iris Odstrcil, Joseph Donovan and Armin Bahl for helpful comments on the manuscript.

## Author Contributions

DG conceived the project with the help of FE. DG conducted the experiments and analyzed the data. CR provided the fish strains used and advised on experiment execution. FE advised on experiments and data analysis. DG, with the help of MH, wrote the manuscript. DG, MH, CR and FE edited the manuscript.

## Methods

### Fish rearing

5-7 days post fertilization (dpf) male and female zebrafish larvae (*Danio rerio*, Hamilton, 1822) were bred in a 14:10 light/dark cycle at 28°C in 10 cm dishes. The strains used were Tg(ath5:SynaptophysinGCaMP6s,isl3:Synaptophysin-GCaMP6s) and Tg(HuC:GCaMP6s), both in a *nac-/-* background. The former expresses GCaMP6s at the axonal terminals of most RGCs in the larval zebrafish, and the latter expresses the same calcium indicator in the soma of most neurons across the whole brain. All animal protocols were in accordance with NIH guidelines and the Harvard University IACUC.

### Stimulus presentation

The larvae were embedded in 1.8% agarose (UltraPure Low Melting Point Agarose, 16520-100, Invitrogen) in a 5 cm plastic petri dish that was then filled with filtered facility water. The stimuli were presented using a custom-built four channel projector. Briefly, two Lightcrafter projectors (Lightcrafter Evaluation Module, Texas Instruments, Dallas, TX, USA) were stacked on top of each other. One of them was modified to support projection of LEDs centered at 606, 463 and 397 nm by changing the dichroic mirrors in the light engine. The second projector was stripped of its dichroic mirrors and the focusing lens was replaced by one supporting UV transmission (354330-A, f=3.1 mm, NA = 0.68, Unmounted Geltech Aspheric Lens, Thorlabs Inc., Newton, NJ, USA). Then, a UV LED (365nm, Mouser Electronics, Mansfield, TX, USA) was mounted at the entrance port of the former red LED. The two projectors were coupled using a flat mirror and a dichroic mirror (PF20-03-F01, Thorlabs Inc.) and their projections were aligned underneath the animal. All stimuli were synthesized by custom software written in LabVIEW (National Instruments, Woburn, MA, USA).

### Brain imaging

Two photon calcium imaging was performed using a custom built point scanning two-photon microscope. Briefly, a Mai-Tai femtosecond laser tuned to 950 nm was passed through a computer controlled half-wave plate and a polarization sensitive prism. After the prism, the beam was expanded to 5 mm and delivered to the center of a set of scanning galvanometric mirrors (Cambridge Technology, Cambridge, MA). The objective used was an Olympus 20X XLUMPLFLN-W water immersion objective (Olympus Corporation, Shinjuku, Tokyo, Japan). Light collected from the sample was relayed to a dichroic mirror that then diverted it to a gateable photomultiplier tube (H11526, Hamamatsu Photonics K.K.), after bandpassing by a filter (Chroma, Bellows Falls, VT). Additionally in the path, there was a Hitachi camera focused on the sample plane to allow for rough sample positioning. The microscope control software was written in LabVIEW (National Instruments, Woburn, MA, USA). The entire system was synchronized so that the stimulus was presented only during the turn around and fly-back periods of the mirror, at which point the PMT was gated off, therefore preventing direct exposure from the projector light. This is important as green light stimulation travels unimpeded through the optical path to the PMT given the emission wavelength of GCaMP. Images were acquired at ∼2 frames per second, at a resolution of 320×320 pixels. Each trial was 40 seconds in length with 10 seconds of adaptation, 20 seconds of stimulus presentation and 10 seconds of rest before the next trial. All stimuli were shown in triplicate to each animal before moving onto the next z section and repeating the whole process. The stimuli presented for Figures 1-3 were full field, single LED intensity oscillations following a sinusoidal temporal structure. During the rest before and after the stimulus, the intensity of the selected LED was left constant and at medium level. For Figure 4, the stimuli were of 3 types: a full field checkerboard with either red or UV in half of the squares and black in the others, oscillating sinusoidally in intensity during the stimulation period and remaining static during the rest periods. The second stimulus was a moving grating alternating the red or UV light with black stripes. The grating was static during rest periods and moved from caudal to nasal during the stimulation period at a spatial frequency of 1cm/cycle and a temporal frequency of 1 cm/s. The third stimulus was a full field red or UV light that suddenly reduced in intensity to 0 for a single frame (900 ms) 20s after the start of the stimulus, and then returned to its original intensity for another 20s.

### Data pre-processing

The raw data followed a pre-processing pipeline that has been described previously [4]. Briefly, the raw imaging data was imported into Matlab (Mathworks, Natick, MA, USA), and all frames within a z section were aligned based on their cross correlation from time point to time point. Following alignment, the ΔF/F was calculated for each stimulus repetition, and the repetitions were collapsed into an average. With the repetitions condensed, all the frames corresponding to one z plane were concatenated and the correlation of each voxel with its 8 neighbors within a plane was calculated across time. An iterative algorithm was used to find the highest correlation value in each z plane and then start grouping it with its neighbors based on a correlation threshold, up to a size threshold or until there were no more neighbors fulfilling the correlation threshold. This was repeated for each high correlation value to yield several groups of voxels. These small groups of voxels, termed ROIs, were then used as the unit to generate each one of the calcium traces used in the study. To obtain each trace, the traces from each voxel in an ROI were averaged together. Finally, signal to noise ratio was approximated as described in Baden et al. [22] by taking the ratio of the averaged standard deviation across repetitions for each trace, and the standard deviation of the average. Then traces below the 25^th^ percentile were excluded from the analysis (around 10% of the traces).

### Calculation of cone gains

This first approximation was calculated assuming linearity in the signal summation from different cone types following Equation 1.

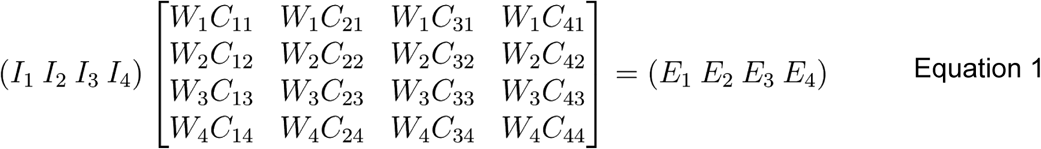

Namely, that a linear combination of the contributions from each cone type (*C*_*ij*_), weighted by constant factors considering the cone and LED spectra (*W*_*i*_), is able to explain the observed responses (*E*_*i*_) as a function of LED intensity (*I*_*i*_). To perform this analysis, the Fourier transform of the traces was extracted to obtain the power at the frequency of stimulation (0.125 Hz). This power was used as the value for the response to each LED. Then, an interaction matrix was constructed, containing the expected excitation of each cone type based on the cone spectra, the LED spectra and the power of the LED. The key element is that, for each stimulus, each LED is turned on on its own, and hence the entire relationship reduces to a four-equation, four-unknown system that can be solved exactly (Equation 2). This will yield a set of cone “weights” for each trace.

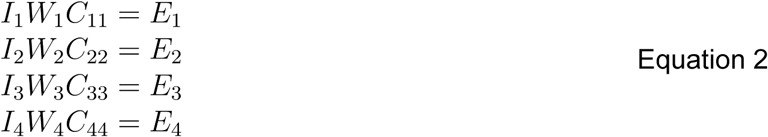

### Gain pattern classification

For identifying the color input patterns into the ROIs we evaluated in this study, we simplified the cone gain patterns analogous to the procedure described in Zhou et al. [8]. Namely, we set the gains within the lowest percentile of average response (5^th^ in our case) to 0, and then any gain with a positive value was assigned a 1 and every gain with a negative value was assigned a −1. Then we determined the unique patterns present in the data and counted the occurrences of each. For plotting, we used the same color convention as described in the reference above, namely, black signifies Off, white signifies 0 and any given color signifies On.

### Registration

The registration procedure was a modification from the one described in Randlett et al. [21]. Briefly, the average stacks of images for each animal and experiment were gaussian blurred and converted to nrrd format, including the metadata for pixel size, and oriented the same as the reference brain (nasal up). Then, the software CMTK (Computational Morphometry Toolkit, http://nitrc.org/projects/cmtk) was utilized to register them to a reference brain, which varied depending on the dataset. The reference brain was cut so as to better match the imaged volume, and hence facilitate the registration process. For the registration itself we used the munger wrapper for CMTK with the following command line parameters: -a -r 01 -l a -v -T 8 -X 52 -C 4 -G 5 -A ‘--accuracy 0.4’ for the isl2 reference brain and (isl3+ath5)::synaptophysin-GCaMP6s data, and -a -r 01 -l a -v -T 8 -X 52 -C 8 -G 20 -A ‘--accuracy 0.4’ for the HuC reference brain and corresponding HuC::GCaMP6s data. Once registered, we loaded the affine matrix into Matlab and applied the registration parameters to the raw calcium data. We translated these from the cut reference brains to the original, full-size reference brains, so as to make them compatible with the full Z Brain Atlas. To produce the maps displayed in the manuscript, after reformatting of the ROIs, we took the average across the stimulation period for each ROI. We then calculated the 75^th^ percentile across all ROIs for each stimulus, and kept only the ROIs above this threshold. After that, we spatially added the signal from all ROIs that crossed the threshold and superimposed this signal on the reference brain stack after normalization of both. Finally, we took the maximum intensity projection in z to generate the final map.

### Similarity index

To obtain a measure of the overlap between the anatomical location of the responding ROIs for each stimulus, we again thresholded the ROIs based on the top 75^th^ percentile of response. Then, we applied a gaussian blurring filter to the position of each remaining ROI to account for the inaccuracies in registration, and next computed the overlap between the ROI sets from each stimulus when positioned in the reference brain on a trial by trial basis. The overlap was quantified as the number of voxels that coincided between the ROIs from each stimulus, without taking intensity into account as this is accounted for by the percentile threshold. To normalize the cross stimulus overlap, we quantified the overlap between single trials of the same stimulus, and then divided the cross-stimulus overlap by the average of the within stimulus overlap for the pair of stimuli in question, thus computing a similarity index that is then plotted in Figures 1K and 4G for their respective datasets.

### Dimensionality reduction and clustering

Given the high dimensionality of the data, the traces were processed using the method described in Baden et al. [22]. Sparse principal component analysis (via the SpaSM Matlab package by Sjöstrand et al. [37]) was used to reduce the dimensionality of each trace. In particular, 4 sPCs were used per stimulus, each one with 10 active bins. To find the ideal number of clusters, the analysis was performed with several cluster numbers and then the one with the minimum Bayesian Information Criterion value was selected. This criterion balances the increase in fit from a stronger model with a penalty to the complexity of the model. The entire process was performed using the Expectation Maximization algorithm in Matlab to fit a Gaussian Mixture Model to the data.

### UMAP

Uniform Manifold Approximation and Projection analysis was performed on the stimulus period of the neural data at the population level. This was done separately for the RGC-RA 4 color dataset, AF10-Tectum 4 color dataset and AF10-Tectum 2 color dataset. We used the implementation for Matlab written by Meehan, Meehan and Moore [38]. The calcium data was projected into PCA space using the sparse principal components used for clustering, normalized, and then processed with the UMAP software. The resulting projection was colored depending on the figure. For Figure 2E, we colored each data point based on its region of origin. or the cluster they belong to. For Figures 3B and 4H we calculated the 75^th^ percentile of the responses and thresholded the ROIs based on this number per stimulus.

### SVM color and intensity classification

To perform the Support Vector Machine classification, the data was supplied to the fitcecoc function in Matlab, which trains a series of binary SVM classifiers to perform multi-class classification. The structure of the binary classifiers is a one-vs-all arrangement, where each binary classifier is trained to separate one category from all the rest (which comprise a single “negative” category together). Then the points that score the highest for a given category are assigned to that one. The classifier for Figure 2G was trained to separate between the four sinusoidal stimuli (i.e. four categories) on a per region basis. We trained one classifier per fish, using only the stimulus period binned to four bins and leave-one-out cross validation. The MATLAB function kfoldpredict was used to obtain the performance for classifying the left out sample in each separate classifier and these were averaged to obtain the classification per fish. Performance of the classifier was assessed as the average percentage of the traces across the diagonal of the confusion matrix (diagram comparing the delivered vs predicted stimulus), either on a per row basis (per stimulus performance) or averaging the whole diagonal (overall performance). The performances of all fish were then averaged to obtain an overall performance and the whole process was repeated ten times. The control classification used the same process, but the labels for the four stimuli were pseudorandomized. For the intensity classifier in Supplementary Figure 2H, we utilized the same procedure outlined above, but instead of using one label per stimulus, we used five levels per stimulus, one for each of the intensity levels of the LED during that stimulus. For the classifier in Figure 3H, we also used the same procedure described above, but we used only the first or the second half of the stimulation period for training, and we binned five time bins into one instead of ten as above. For the classifier in Figure 4J we used an analogous procedure as for Figure 2G, but we only used the pair of red and UV stimuli corresponding to any given pattern and trained separate binary classifiers. For the classifier in Figure 4K instead, we used ten time bins from the pre-stimulus period and the ten time bins in the middle of the stimulation period, as the goal was to classify between stimulus vs no stimulus. This was done also for each pair of red and UV corresponding to a single pattern at a time.

### Cluster to cluster regression

As an alternative way to assess the similarity between the responses in each region, we calculated the average of each cluster and region, and then used cross-validated linear regression to fit the responses of one region with another. This was done for all the possible pairings of regions. Then, for each pair, the losses from the fits to each cluster were averaged and saved in a matrix. Finally, we plotted the results as 1 minus the average loss, as we wanted a measure of similarity.

### Correlation analysis

Four different types of correlation were calculated, all using Pearson’s correlation coefficient. For calculation of the correlation matrices in Figure 3E, the data sets were reshaped so that all of the responses for a given stimulus are concatenated across neurons, and separated by stimulus. Then a correlation matrix was calculated based on this input so that the result had the dimensions of the number of stimuli. This was done with both the four sinusoidal stimuli data set and the red/UV pattern data set. To calculate the correlations shown in Figure 3E, the cluster averages from the responses for the different stimuli within a brain region (AF10 or tectum) were correlated against each other, yielding a pairwise correlation matrix of color comparisons for that brain region. For the correlation over time matrix in Figure 3F-G, the traces were separated by stimulus, and the individual response vectors for each time point were correlated against each other, yielding a single correlation coefficient per stimulus. For calculation of the stimulus-to-stimulus correlation over time in Supplementary Figure 3D, the traces were again concatenated across neurons, but they were kept separate for each time point. Then, the traces for each stimulus were correlated with each other at each time point, yielding the trajectories displayed. All four of the above were calculated individually per animal and then averaged. For calculation of the fish to fish correlations, the data from all the animals in the tectal data set was clustered together, then the traces from each animal were averaged based on their belonging to a cluster. These averages were then correlated across animals and an average correlation per individual was calculated.

### PCA-CCA

The Principal Component Analysis was performed for each animal separately, using the neural response data for the stimulus period only. As outlined in Gallego et al. [24], although PCA realigns the space the data is in uniquely for that data (i.e. set of neurons/axonal terminals), if the underlying variance-driving dimensions are similar, or in other words if the sets of signals from different animals lie in the same manifold of activity, one can use Canonical Correlation Analysis to find a common subspace that maximizes the alignment between the PCA reconstructions from different sets of ROIs. We therefore utilized this technique to align the PCA decompositions of the stimulus responses for the AF10 and tectum from different animals. We only took the animals that had at least three dimensions containing 50% of the variance or higher. We then averaged these trajectories to generate the displays shown in Figure 3I. These aligned trajectories were also used to compute the results in Figure 3K.

## References

1. Raymond, P.A., Barthel, L.K., and Curran, G.A. (1995). Developmental patterning of rod and cone photoreceptors in embryonic zebrafish. J. Comp. Neurol. 359, 537–550.

2. Robinson, J., Schmitt, E.A., Harosi, F.I., Reece, R.J., and Dowling, J.E. (1993). Zebrafish ultraviolet visual pigment: absorption spectrum, sequence, and localization. Proceedings of the National Academy of Sciences 90, 6009–6012. Available at: http://dx.doi.org/10.1073/pnas.90.13.6009.

3. Spence, R., Gerlach, G., Lawrence, C., and Smith, C. (2008). The behaviour and ecology of the zebrafish, Danio rerio. Biol. Rev. Camb. Philos. Soc. 83, 13–34.

4. Portugues, R., Feierstein, C.E., Engert, F., and Orger, M.B. (2014). Whole-brain activity maps reveal stereotyped, distributed networks for visuomotor behavior. Neuron 81, 1328–1343.

5. Ahrens, M.B., Orger, M.B., Robson, D.N., Li, J.M., and Keller, P.J. (2013). Whole-brain functional imaging at cellular resolution using light-sheet microscopy. Nat. Methods 10, 413–420.

6. Zimmermann, M.J.Y., Nevala, N.E., Yoshimatsu, T., Osorio, D., Nilsson, D.-E., Berens, P., and Baden, T. (2018). Zebrafish Differentially Process Colour Across Visual Space to Match Natural Scenes. Current Biology. Available at: http://dx.doi.org/10.2139/ssrn.3155573.

7. Connaughton, V.P., and Nelson, R. (2010). Spectral Responses in Zebrafish Horizontal Cells Include a Tetraphasic Response and a Novel UV-Dominated Triphasic Response. Journal of Neurophysiology 104, 2407–2422. Available at: http://dx.doi.org/10.1152/jn.00644.2009.

8. Zhou, M., Bear, J., Roberts, P.A., Janiak, F.K., Semmelhack, J., Yoshimatsu, T., and Baden, T. (2020). Zebrafish Retinal Ganglion Cells Asymmetrically Encode Spectral and Temporal Information across Visual Space. Curr. Biol. Available at: http://dx.doi.org/10.1016/j.cub.2020.05.055.

9. Ma, M., Kler, S., and Pan, Y.A. (2019). Structural Neural Connectivity Analysis in Zebrafish With Restricted Anterograde Transneuronal Viral Labeling and Quantitative Brain Mapping. Front. Neural Circuits 13, 85.

10. Wulliman, M.F., Rupp, B., and Reichert, H. (2012). Neuroanatomy of the Zebrafish Brain: A Topological Atlas (Birkhäuser).

11. Robles, E., Laurell, E., and Baier, H. (2014). The retinal projectome reveals brain-area-specific visual representations generated by ganglion cell diversity. Curr. Biol. 24, 2085–2096.

12. Burrill, J.D., and Easter, S.S., Jr (1994). Development of the retinofugal projections in the embryonic and larval zebrafish (Brachydanio rerio). J. Comp. Neurol. 346, 583–600.

13. Niessing, J., and Friedrich, R.W. (2010). Olfactory pattern classification by discrete neuronal network states. Nature 465, 47–52.

14. Johnson, K.O. (2000). Neural Coding. Neuron 26, 563–566.

15. Estévez, O., and Spekreijse, H. (1982). The “silent substitution” method in visual research. Vision Res. 22, 681–691.

16. Chen, T.-W., Wardill, T.J., Sun, Y., Pulver, S.R., Renninger, S.L., Baohan, A., Schreiter, E.R., Kerr, R.A., Orger, M.B., Jayaraman, V., et al. (2013). Ultrasensitive fluorescent proteins for imaging neuronal activity. Nature 499, 295–300.

17. Mitchell, D.E., and Rushton, W.A. (1971). Visual pigments in dichromats. Vision Res. 11, 1033–1043.

18. Risner, M.L., Lemerise, E., Vukmanic, E.V., and Moore, A. (2006). Behavioral spectral sensitivity of the zebrafish (Danio rerio). Vision Res. 46, 2625–2635.

19. McDowell, A.L., Dixon, L.J., Houchins, J.D., and Bilotta, J. (2004). Visual processing of the zebrafish optic tectum before and after optic nerve damage. Vis. Neurosci. 21, 97–106.

20. Cameron, D.A. (2002). Mapping absorbance spectra, cone fractions, and neuronal mechanisms to photopic spectral sensitivity in the zebrafish. Vis. Neurosci. 19, 365–372.

21. Randlett, O., Wee, C.L., Naumann, E.A., Nnaemeka, O., Schoppik, D., Fitzgerald, J.E., Portugues, R., Lacoste, A.M.B., Riegler, C., Engert, F., et al. (2015). Whole-brain activity mapping onto a zebrafish brain atlas. Nat. Methods 12, 1039–1046.

22. Baden, T., Berens, P., Franke, K., Román Rosón, M., Bethge, M., and Euler, T. (2016). The functional diversity of retinal ganglion cells in the mouse. Nature 529, 345–350.

23. Becht, E., McInnes, L., Healy, J., Dutertre, C.-A., Kwok, I.W.H., Ng, L.G., Ginhoux, F., and Newell, E.W. (2018). Dimensionality reduction for visualizing single-cell data using UMAP. Nat. Biotechnol. Available at: http://dx.doi.org/10.1038/nbt.4314.

24. Gallego, J.A., Perich, M.G., Chowdhury, R.H., Solla, S.A., and Miller, L.E. (2020). Long-term stability of cortical population dynamics underlying consistent behavior. Nat. Neurosci. 23, 260–270.

25. Livingstone, M. (2008). Vision and Art: The Biology of Seeing (Harry N. Abrams).

26. Baden, T., and Osorio, D. (2019). The Retinal Basis of Vertebrate Color Vision. Annu Rev Vis Sci 5, 177–200.

27. Grama, A., and Engert, F. (2012). Direction selectivity in the larval zebrafish tectum is mediated by asymmetric inhibition. Front. Neural Circuits 6, 59.

28. Nikolaou, N., Lowe, A.S., Walker, A.S., Abbas, F., Hunter, P.R., Thompson, I.D., and Meyer, M.P. (2012). Parametric functional maps of visual inputs to the tectum. Neuron 76, 317–324.

29. Baumann, O., Borra, R.J., Bower, J.M., Cullen, K.E., Habas, C., Ivry, R.B., Leggio, M., Mattingley, J.B., Molinari, M., Moulton, E.A., et al. (2015). Consensus paper: the role of the cerebellum in perceptual processes. Cerebellum 14, 197–220.

30. Bird, C.M., Berens, S.C., Horner, A.J., and Franklin, A. (2014). Categorical encoding of color in the brain. Proc. Natl. Acad. Sci. U. S. A. 111, 4590–4595.

31. Namboodiri, V.M.K., Rodriguez-Romaguera, J., and Stuber, G.D. (2016). The habenula. Curr. Biol. 26, R873–R877.

32. Semmelhack, J.L., Donovan, J.C., Thiele, T.R., Kuehn, E., Laurell, E., and Baier, H. (2014). A dedicated visual pathway for prey detection in larval zebrafish. Elife 3. Available at: http://dx.doi.org/10.7554/eLife.04878.

33. Zhang, B.-B., Yao, Y.-Y., Zhang, H.-F., Kawakami, K., and Du, J.-L. (2017). Left Habenula Mediates Light-Preference Behavior in Zebrafish via an Asymmetrical Visual Pathway. Neuron 93, 914–928.e4.

34. Knogler, L.D., Markov, D.A., Dragomir, E.I., Štih, V., and Portugues, R. (2017). Sensorimotor Representations in Cerebellar Granule Cells in Larval Zebrafish Are Dense, Spatially Organized, and Non-temporally Patterned. Curr. Biol. 27, 1288–1302.

35. Halpern, M.E., Liang, J.O., and Gamse, J.T. (2003). Leaning to the left: laterality in the zebrafish forebrain. Trends Neurosci. 26, 308–313.

36. Conway, B.R., Eskew, R.T., Jr, Martin, P.R., and Stockman, A. (2018). A tour of contemporary color vision research. Vision Res. 151, 2–6.

37. Sjöstrand, K., Clemmensen, L.H., Larsen, R., Einarsson, G., and Ersbøll, B. (2018). SpaSM: A MATLAB Toolbox for Sparse Statistical Modeling. Journal of Statistical Software 84. Available at: http://dx.doi.org/10.18637/jss.v084.i10.

38. Meehan, C., Meehan, S., and Moore, W. (2020). Uniform Manifold Approximation and Projection (UMAP), MATLAB Central File Exchange Available at: https://www.mathworks.com/matlabcentral/fileexchange/71902).

