## Supplementary Material for "Colors everywhere: enhanced chromatic processing across the first visual synapse in the zebrafish central brain"

### Supplementary Information

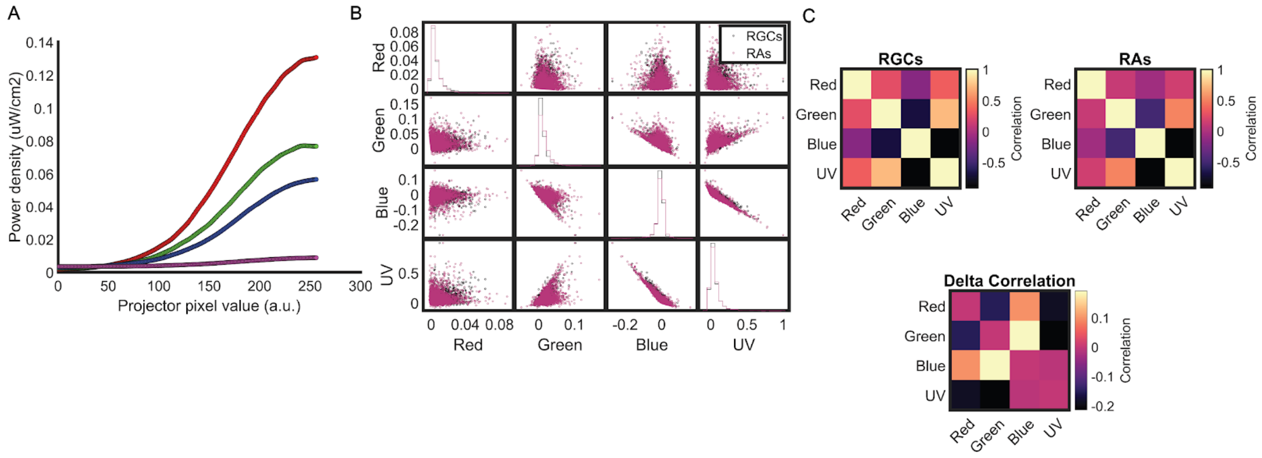


**Supplementary Figure 1 - Related to Figure 1.** A: pixel intensity to LED power curves for the projector used. B: correlation of the cone gains for both the RGCs and RAs on a single ROI basis. The scatter plots relate the values for the given cone types, and the histograms show the distribution of values for that cone type. C: (top) overall correlation between the cone gains for RGCs and RAs. (bottom) delta cone gains correlation for each cone type. Blue and UV concentrate the lowest correlation.

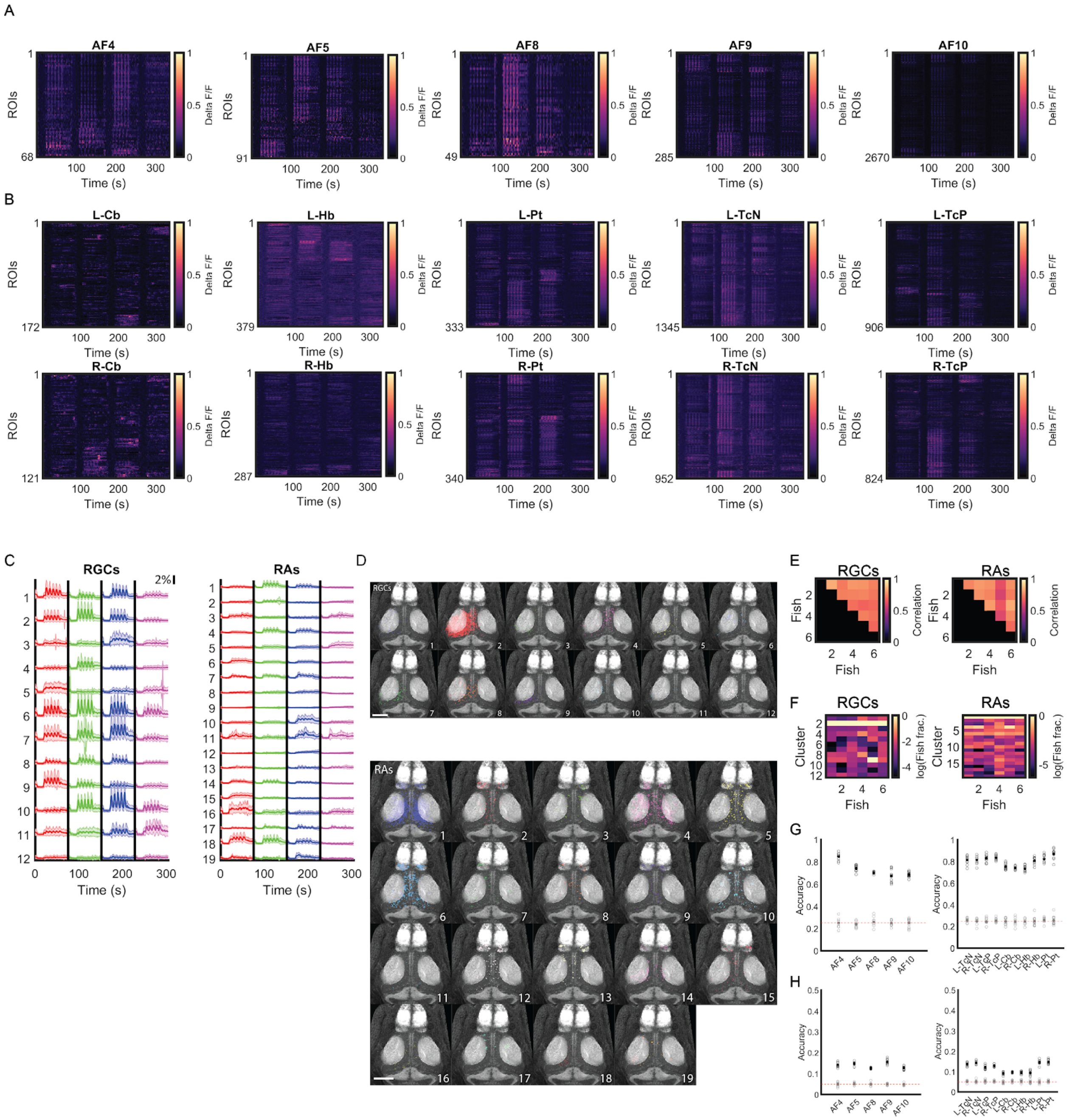


**Supplementary Figure 2 - Related to Figure 2.** A: calcium traces corresponding to the indicated brain regions within the RGC termination fields. Intensity is delta F/F. B: same as A for the RA brain regions. C: cluster average traces for the RGC and RA brain regions. D: anatomical location of the clusters indicated in C for the RGC and RA brain regions. E: correlation between cluster averages from each fish in the RGC and RA datasets. F: proportion of every cluster in each individual fish for RGCs and RAs. G: average classification performance from an SVM classifier trained to distinguish between the four color stimuli for each region (black circles, average in filled circle) compared to shuffled controls (gray circles, average in filled circle). H: SVM classifier performance per region when the classifier was trained to discern between the individual intensity levels across the stimuli, compared to shuffled controls. Labelling scheme is the same as G. Scale bars in C are 90 µm. AF: arborization field, TcN: tectal neuropil, TcP: tectal periventricular region, Pt: pretectum, Hb: habenula, Cb: cerebellum, R- L-; right or left hemisphere.

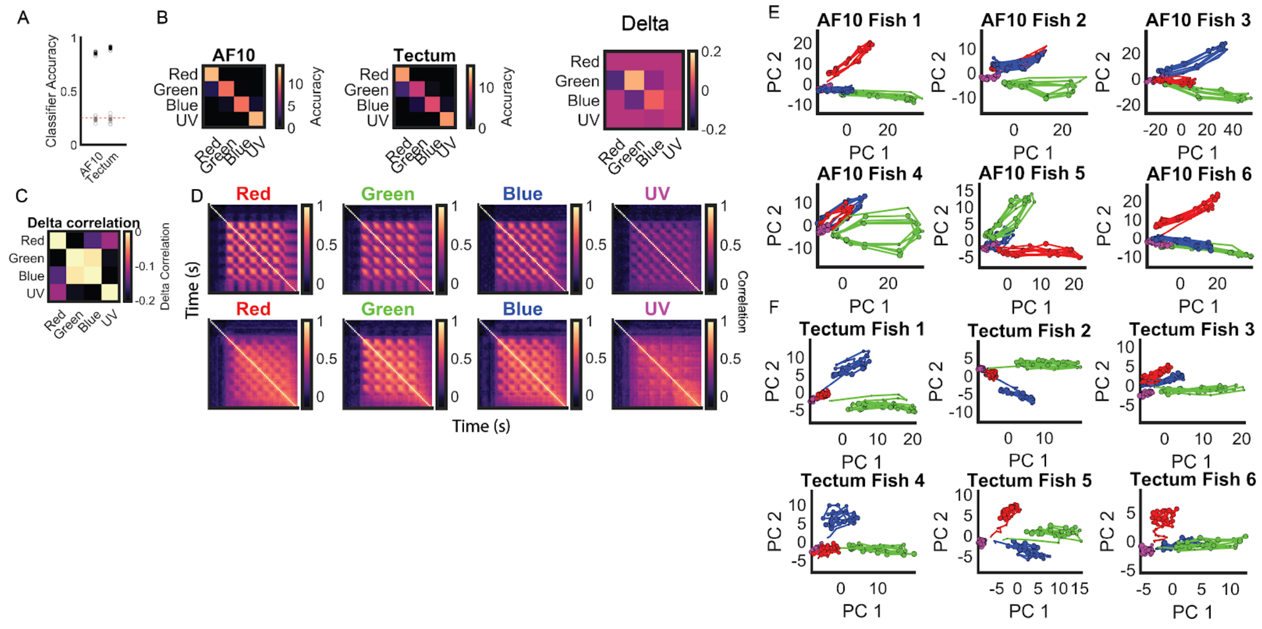


**Supplementary Figure 3 - Related to Figure 3.** A: average classification performance (black circles are individual training repetitions with the average in the filled black circle) of an SVM trained to discern between the four color stimuli compared to shuffled controls (gray circles are individual repetitions and the filled gray circle is the average). B: average confusion matrix of ten repetitions of the classifier trained in A for both AF10 and tectum (left and middle). Delta confusion matrix of the previous classifiers to highlight the differences (right). C: delta correlation matrix when comparing the correlation matrices from both datasets shown in Fig. 3D. D: correlation over time for each stimulus response in both AF10 (top) and tectum (bottom). E: individual fish PCA projections of the neural activity during the stimulation period for AF10. F: same as in E for the tectum.

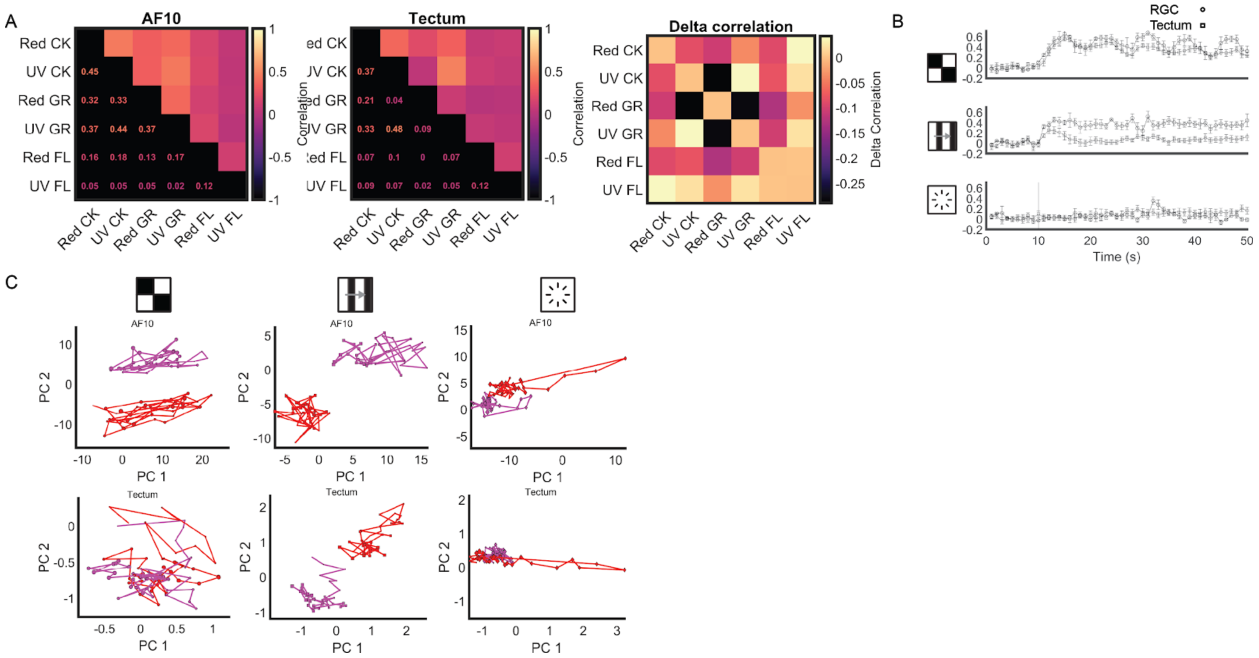


**Supplementary Figure 4 - Related to Figure 4.** A: correlation matrices for the responses evoked by each of the six stimuli in the set for AF10 (left) and tectum (middle). Delta correlation matrix for the two previous matrices to highlight their differences (right). Overall the level of correlation is high. B: correlation over time for the responses evoked by the different patterns irrespective of color in AF10 and tectum. The patterns do not differentiate well. C: CCA-aligned average PCA trajectories for each one of the six stimuli in AF10 and tectum. Colors remain mostly separated.
